# A high-throughput flow cytometry assay to evaluate CatSper-mediated Ca^2+^ influx triggered by high-K^+^/high-pH stimulation in mouse sperm

**DOI:** 10.64898/2026.01.05.697741

**Authors:** Lucila R. Gomez-Olivieri, Liza J. Schiavi-Ehrenhaus, Martina Jabloñski, Mariano G. Buffone, Guillermina M. Luque

## Abstract

The sperm-specific Ca^2+^ channel CatSper is essential for male fertility, as it mediates Ca^2+^ influx required for sperm hyperactivation. Because of its restricted expression in sperm and crucial role in fertilization, CatSper represents a promising target for the development of non-hormonal male contraceptives. However, its structural complexity and the absence of functional heterologous expression systems have limited inhibitor discovery. Here, we developed a high-throughput flow cytometry assay to identify compounds capable of blocking CatSper activity in mammalian sperm. The method relies on the stimulation of CatSper by exposing live mouse sperm to a high-K^+^/high-pH solution (K8.6), which induces membrane depolarization and intracellular alkalinization, triggering Ca^2+^ entry through CatSper. Changes in intracellular Ca^2+^ levels were monitored using the fluorescent indicator Fluo-4. To increase throughput and minimize variability, this assay was combined with fluorescent cell barcoding, which assigns each sperm sample a unique fluorescent signature, enabling the simultaneous acquisition of multiple conditions within a single tube. This integration greatly reduces acquisition time and reagent consumption, thereby facilitating large-scale screening experiments. Moreover, we implemented a compound pooling strategy to further improve assay efficiency, allowing the rapid identification of active compounds within a library. The assay was validated using CatSper1 knockout sperm and known pharmacological inhibitors, confirming that both genetic deletion and pharmacological blockade of CatSper abolish the K8.6-induced Ca^2+^ response. Together, these results establish a robust, scalable, and high-throughput screening platform for identifying novel CatSper blockers and potential male contraceptive candidates.

## Introduction

The sperm-specific Ca^2+^ channel CatSper is the principal entity for Ca^2+^ entry and plays an essential role in the development of hyperactivated motility, a vigorous and asymmetric flagellar movement required for fertilization [1–4]. Genetic deletion of CatSper subunits in mice, as well as loss-of-function mutations in humans, result in sperm unable to hyperactivate and, consequently, in male infertility [4–7].

CatSper is one of the most complex ion channels characterized to date. It consists of four pore-forming α subunits (CatSper1–4) [8,9] accompanied by a large number of auxiliary subunits, including: CatSperβ, γ, δ, ε, ζ, η, τ, EFCAB9, SLCO6C1, and TMEM249, among others [10–17]. These proteins assemble in the plasma membrane of the principal piece, forming four longitudinal nanodomains of Ca^2+^ signaling along the flagellum [13,14,18]. This unique structural organization makes CatSper extraordinarily challenging to study, as its heterologous expression has not been achieved and *in vitro* biochemical analyses remain limited.

Soon after the discovery of CatSper1 α subunit [1], the first electrophysiological recordings of mouse sperm were achieved using the whole-cell patch-clamp technique [19], and later adapted to human sperm [20]. Despite its major impact, the sperm patch-clamp technique is technically demanding and performed by only a few laboratories worldwide, as it requires specialized equipment and a unique personal expertise [9]. Consequently, although it is the only direct and gold standard method for assessing CatSper function, its application is limited to highly specialized settings.

To overcome these limitations, indirect approaches have been implemented to evaluate CatSper opening. Measuring changes in the sperm membrane potential (Em) after chelating extracellular Ca^2+^ with EGTA and using the fluorescent probe DiSC3(5) makes it possible to quantify CatSper-dependent depolarization [19,21,22]. This assay, performed using both spectrofluorometry and flow cytometry, has proven to be a robust and reproducible indirect indicator of CatSper opening [23–30].

Another indirect strategy to assess CatSper activity exploits the complete cessation of sperm motility observed in Ca^2+^-free conditions [4,31] due to the influx of Na^+^ through CatSper and the resulting consumption of sperm ATP [22]. Sperm lacking functional CatSper remain motile under these conditions [4,31]. This assay, recently adapted for human sperm, has proven useful to confirm the dependence of sperm motility on CatSper and to identify patients with defective CatSper function [4].

A further well-established strategy to trigger CatSper activity is to depolarize and alkalinize sperm with a high-K^+^/high-pH medium (K8.6). This treatment shifts the voltage dependence of CatSper toward more negative potentials, due to the rapid intracellular alkalinization, thereby rendering the channel more prone to open upon membrane depolarization [9,19]. As a result, K8.6 elicits a rapid and sustained Ca^2+^ influx through CatSper, as demonstrated in both mouse and human sperm [32,33]. The K8.6-induced increase in intracellular Ca^2+^ concentration ([Ca^2+^]ᵢ) has been used as a functional readout of CatSper activity in various experimental settings, including high-throughput screening assays for identifying CatSper inhibitors [33]. However, current implementations of this approach rely mostly on either bulk fluorescence measurements, such as spectrofluorometry, which provide only the average response of the entire sperm population [33], or single-cell live imaging, which captures individual responses but typically from a limited number of cells [32,34]. Both approaches therefore restrict the ability to detect functional heterogeneity within sperm populations and to correlate CatSper-dependent Ca^2+^ responses with additional physiological parameters. In this study, a flow cytometry-based adaptation of the K8.6-induced CatSper activation assay was validated. This approach enables sensitive, single-cell resolution analysis of CatSper-mediated Ca^2+^ entry in a large number of sperm, allowing discrimination of functional heterogeneity within sperm populations, as well as providing individual fluorescence data and differentiation between live and dead cells. Moreover, the method can be combined with fluorescence cell barcoding (FCB), enabling its integration into multiparametric experimental designs.

## Materials and methods

### Reagents

All reagents are detailed in **Supplementary Table 1**.

### Animals

Mature (10–12-week-old) male hybrid F1 (female BALB/c × male C57BL/6) mice and CatSper1 knockout (KO) [1] mice along with their corresponding heterozygous (HET) C57BL/6 male siblings were used. Animals were group-housed (4–5 per cage) under controlled environmental conditions (23°C, 12 h light/dark cycle; lights on 07:00–19:00 h) with *ad libitum* access to food and water. All experimental procedures were reviewed and approved by the Ethical Committees of the IBYME, Buenos Aires (#32/2021) and were performed in strict accordance with the Guide for the Care and Use of Laboratory Animals approved by the National Institutes of Health (NIH).

### Sperm medium and collection

The non-capacitating medium (NC medium) used in this study was a modified Toyoda–Yokoyama–Hosi (modified TYH) containing 119.3 mM NaCl, 4.7 mM KCl, 1.71 mM CaCl2.2H2O, 1.2 mM KH2PO4, 1.2 mM MgSO4.7H2O, 0.51 mM sodium pyruvate, 5.56 mM glucose, 20 mM HEPES, and 10 μg/mL gentamicin, pH 7.35–7.40. Animals were euthanized and both cauda epididymis were placed in 500 μl of NC medium. After 15 min of incubation at 37°C (swim-out) epididymis were removed.

### Evaluation of CatSper opening by measuring [Ca^2+^]ᵢ changes by flow cytometry

Sperm [Ca^2+^]ᵢ was assessed using Fluo-4 AM as previously described [35]. Briefly, sperm obtained by swim-out were incubated in NC medium containing 1 μM Fluo-4 AM and 0.02% pluronic acid for 20 min at 37°C. Samples were washed and resuspended in NC medium containing either vehicle (DMSO) or CatSper inhibitors (20 μM HC-056456, 10 μM RU1968 or 10 μM 5-(N,N-hexamethylene)-Amiloride (HMA)). Before collecting data, 2 ng/μL propidium iodide (PI) was added to monitor viability. Data were acquired as individual cellular events using a BD FACSCanto II TM cytometer (Biosciences; Becton, Dickinson and Company). Basal [Ca^2+^]ᵢ levels were recorded for 30 s of continuous acquisition. Afterwards, samples were centrifugated at 700 g for 4 min, and NC medium was removed. Immediately, K8.6 medium (135 mM KCl, 5 mM NaCl, 2 mM CaCl2, 1 mM MgCl2, 10 mM glucose, 10 mM lactic acid, 1 mM sodium pyruvate, and 30 mM HEPES, pH 8.60) [36] was added to induced CatSper activation, and acquisition continued for additional 90 s. Doublets were excluded using forward-scatter area (FSC-A) vs. forward-scatter height (FSC-H) dot plots. Fluo-4 fluorescence was acquired using the fluorescein isothiocyanate (FITC; 530/30) detection channel, while PI fluorescence was recorded using the peridinin chlorophyll protein (PerCP; 670 LP) channel. The two indicators had minimal emission overlap, but compensation was still done. Data was analyzed using FlowJo software (V10.0.7), and the fluorescence median was normalized to the basal fluorescence in each condition.

### FCB protocol in combination with the high-K^+^/high-pH assay

FCB was adapted following the protocol previously described [30]. Three fluorescent dyes were used: Fluo-4 AM to monitor [Ca^2+^]ᵢ changes, and CellTrace™ Violet together with CellTrace™ Far Red for barcoding. Eight distinct barcodes were generated using four concentrations of CellTrace™ Violet (0.025, 1.5, 5, and 20 µM) combined with two concentrations of CellTrace™ Far Red (0.001 and 0.4 µM). For each dye, serial dilutions were prepared in NC medium from 5 mM (Violet) and 1 mM (Far Red) stock solutions.

For staining, 35 µL of each FCB dye (35 µL CellTrace™ Violet 4X + 35 µL CellTrace™ Far Red 4X), 35 µL of Fluo-4 AM 4X, and 0.5 µL of vehicle or compound pools (three compounds/well) were combined and mixed with 35 µL of sperm suspension (30–40 × 10⁶/mL; pipette tip immersed to ensure uniform labeling), yielding a final volume of 140.5 µL per well, 1 μM Fluo-4 AM and 0.02% pluronic acid, and a final sperm concentration of 7.5–10 × 10⁶/mL. 96-well plates were incubated for 20 min at 37°C in the dark, covered to prevent evaporation and dye photobleaching. After incubation, 195 µL of NC medium was added to each well. Barcoded samples were then pooled by transferring the entire content of each well into a 15 mL tube containing 5 mL of pre-warmed NC medium to dilute the dyes and compounds and stop the labeling reaction. Cells were washed by centrifugation at 700 g for 15 min at room temperature to remove any unbound dyes. Immediately after centrifugation, 150 µL of the pellet was transferred to a new microcentrifuge tube for acquisition. Flow cytometric data were acquired as previously described. Detection channels were set as follows: Fluo-4 AM, FITC (530/30 nm); CellTrace™ Violet, Pacific Blue™/AmCyan (450/50 nm); CellTrace™ Far Red, APC (660/20 nm); and PI, PerCP (670 LP). For CatSper activation, [Ca^2+^]i was recorded for 30 s under basal conditions, followed by replacement of NC medium with K8.6 medium, and acquisition continued for an additional 90 s. In all cases, doublet exclusion was performed as previously described. Compensation was done by performing the appropriate unmixed barcoded samples compensation controls in each experiment. Data were analyzed using FlowJo software (V10.0.7).

## Statistical analysis

Statistical analyses were performed using the GraphPad Prism 9 software (La Jolla, CA, USA). Data are expressed as the mean ± standard error of the mean (SEM). In all cases, two-way ANOVA for matched data with Sidak’s multiple comparisons test was performed. P < 0.05 was considered statistically significant.

## Results

### The high-K⁺/high-pH assay by flow cytometry is a suitable tool for analyzing CatSper function

The K8.6-induced [Ca^2+^]ᵢ increase assay was optimized for flow cytometry, allowing the rapid acquisition of data from thousands of individual sperm while simultaneously distinguishing live from dead cells and quantifying intracellular fluorescence at single-cell resolution. Sperm [Ca^2+^]ᵢ levels were monitored using the fluorescent Ca^2+^ indicator Fluo-4 together with PI to selectively analyze viable cells, as previously described [35].

Under basal conditions, Fluo-4–loaded sperm analyzed by flow cytometry revealed two distinct populations: one exhibiting low and the other high fluorescence intensity, indicating heterogeneity in basal [Ca^2+^]ᵢ among cells (**Figure 1A–B**), consistent with our previous report [35]. Upon exposure to the K8.6 medium, a clear increase in [Ca^2+^]ᵢ was detected only in sperm expressing functional CatSper (CatSper1 HET), whereas sperm lacking CatSper (CatSper1 KO) failed to respond and remained within the low-fluorescence population (**Figure 1A–C**).

**Figure 1.**
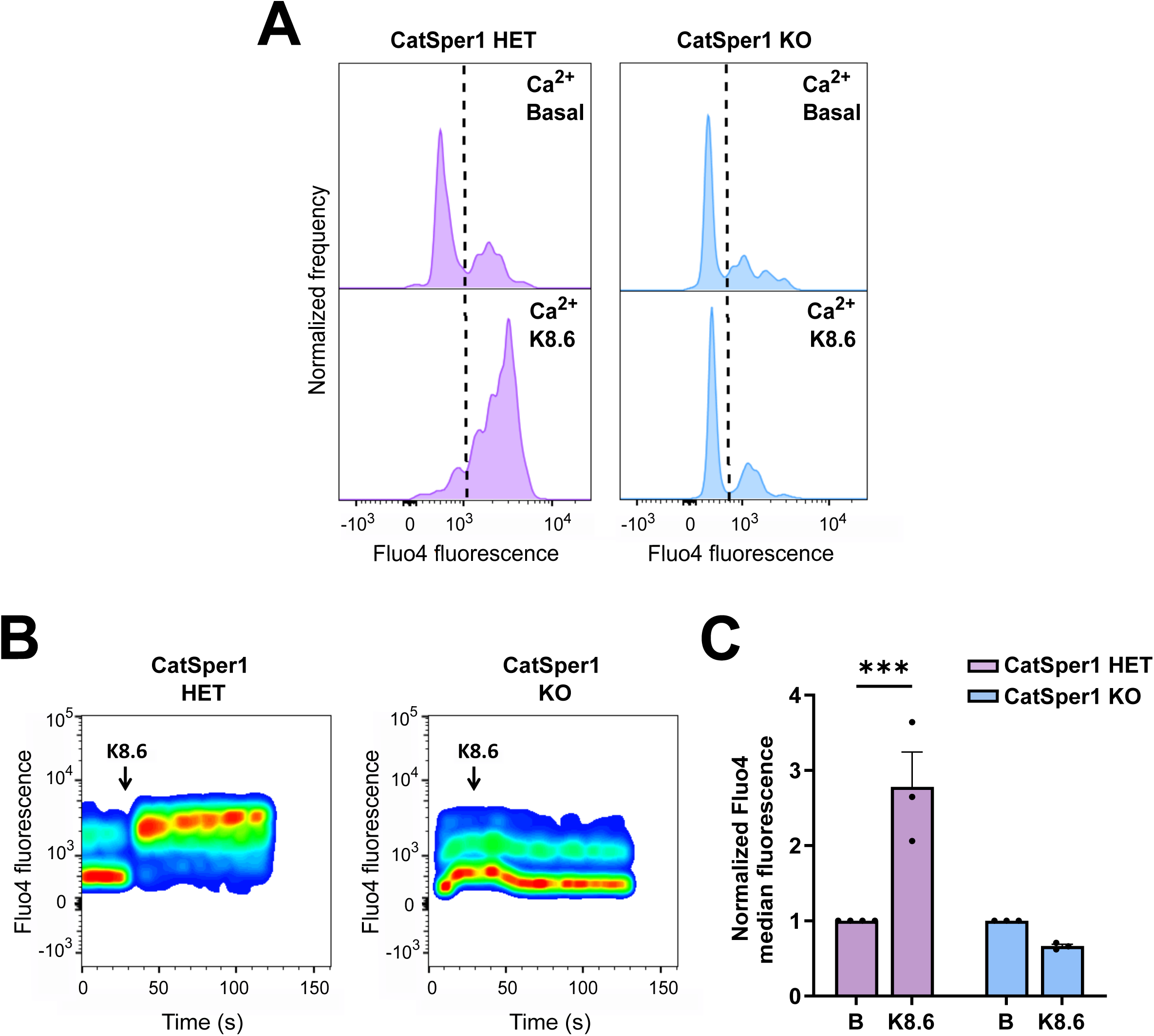
Assessment of CatSper activity by flow cytometry in live mouse sperm. CatSper activity was evaluated in CatSper1 HET and KO sperm by monitoring K8.6-induced [Ca^2+^]ᵢ increases using Fluo-4. **(A)** Representative histograms showing Fluo-4 fluorescence before (basal [Ca^2+^]ᵢ) and after K8.6 stimulation. **(B)** Representative 2D dot plots of Fluo-4 fluorescence over time; K8.6 addition is indicated by a black arrow. The responsive population was defined under the CatSper1 HET condition and used as reference for CatSper1 KO samples. **(C)** Median Fluo-4 fluorescence before (B) and after stimulation (K8.6) normalized to the basal condition (n=3). ***p < 0.001 indicates statistical difference.

As shown in **Figure 1C**, K8.6 treatment produced a significant rise in the normalized median Fluo-4 fluorescence of live CatSper1 HET sperm, but not in CatSper1 KO cells. These results confirm that the flow cytometry–based K8.6 assay provides a robust and sensitive approach for evaluating CatSper-mediated Ca^2+^ entry.

Consistently, exposure of sperm to the CatSper inhibitors HC-056456, RU1968, or HMA [33,37,38] completely suppressed the K8.6-evoked [Ca^2+^]ᵢ elevation (**Figure 2A–B**), resembling CatSper1 KO phenotype.

**Figure 2.**
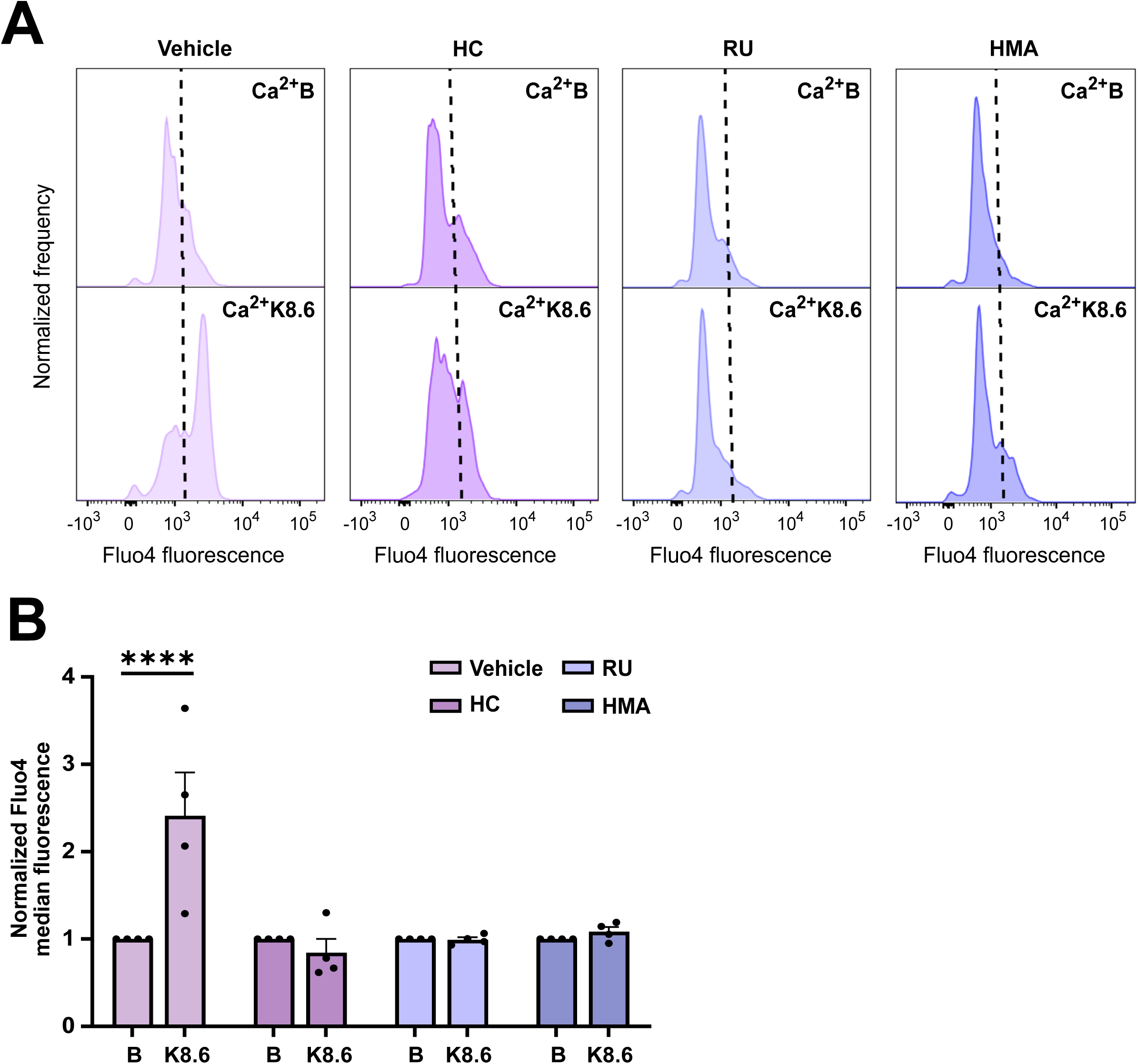
Validation of CatSper activity assessment using CatSper inhibitors. CatSper activity in live sperm was analyzed by flow cytometry after stimulation with K8.6 medium. Sperm were pre-incubated for 20 min with Fluo-4 dye, treated with vehicle (DMSO) or CatSper inhibitors (20 μM HC-056456, 10 μM RU1968, or 10 μM HMA), and [Ca^2+^]ᵢ changes were monitored before and after K8.6 addition. **(A)** Representative histograms of control and inhibitor-treated sperm showing Fluo-4 fluorescence before (basal=B) and after K8.6 stimulation. **(B)** Median Fluo-4 fluorescence before (B) and after K8.6 normalized to the basal condition (n ≥ 3). ****p < 0.0001 indicates statistical difference.

### Development of a high-throughput flow cytometry assay for CatSper screening by combining FCB and the K8.6 activation method

Based on our previous work demonstrating that FCB can be successfully applied to live sperm [30], we adapted this strategy to establish a high-throughput assay for assessing CatSper channel activity. FCB allows the simultaneous acquisition of several samples within a single flow cytometry run, increasing reproducibility while reducing both reagent use and experimental variability [39,40]. Each sample is labeled with a unique combination of fluorescent dyes, generating a distinct spectral “barcode” that can later be identified during data analysis.

To integrate this approach with a CatSper functional assay, we combined FCB with the K8.6-induced Ca^2+^ entry test. Two spectrally compatible dyes, CellTrace™ Far Red and CellTrace™ Violet, were optimized to yield eight clearly distinguishable barcodes that did not interfere with the emission spectra of Fluo-4 (Ca^2+^ indicator) or PI (used for viability assessment).

To make the screening process more efficient, we implemented a compound pooling strategy in which three compounds were tested together per well. This design substantially reduces the number of assays needed, as non-active mixtures can be rapidly discarded based on the absence of CatSper inhibition, following strategies described for other screening platforms [41].

Figure 3 shows a representative schematic (not based on experimental data) illustrating the overall workflow. Sperm were co-incubated with Fluo-4, the FCB dyes, and compound mixtures under NC conditions. Labeled samples were then combined into a single tube, washed, and stained with PI to exclude dead cells. Fluorescence was recorded before and after exposure to K8.6 medium to activate CatSper channels. Data analysis using FlowJo software revealed distinct sperm populations corresponding to each barcode in two-dimensional plots of Violet vs. Far Red fluorescence, allowing the comparison of Ca^2+^ responses across all experimental conditions (Figure 3**, step 7**).

**Figure 3.**
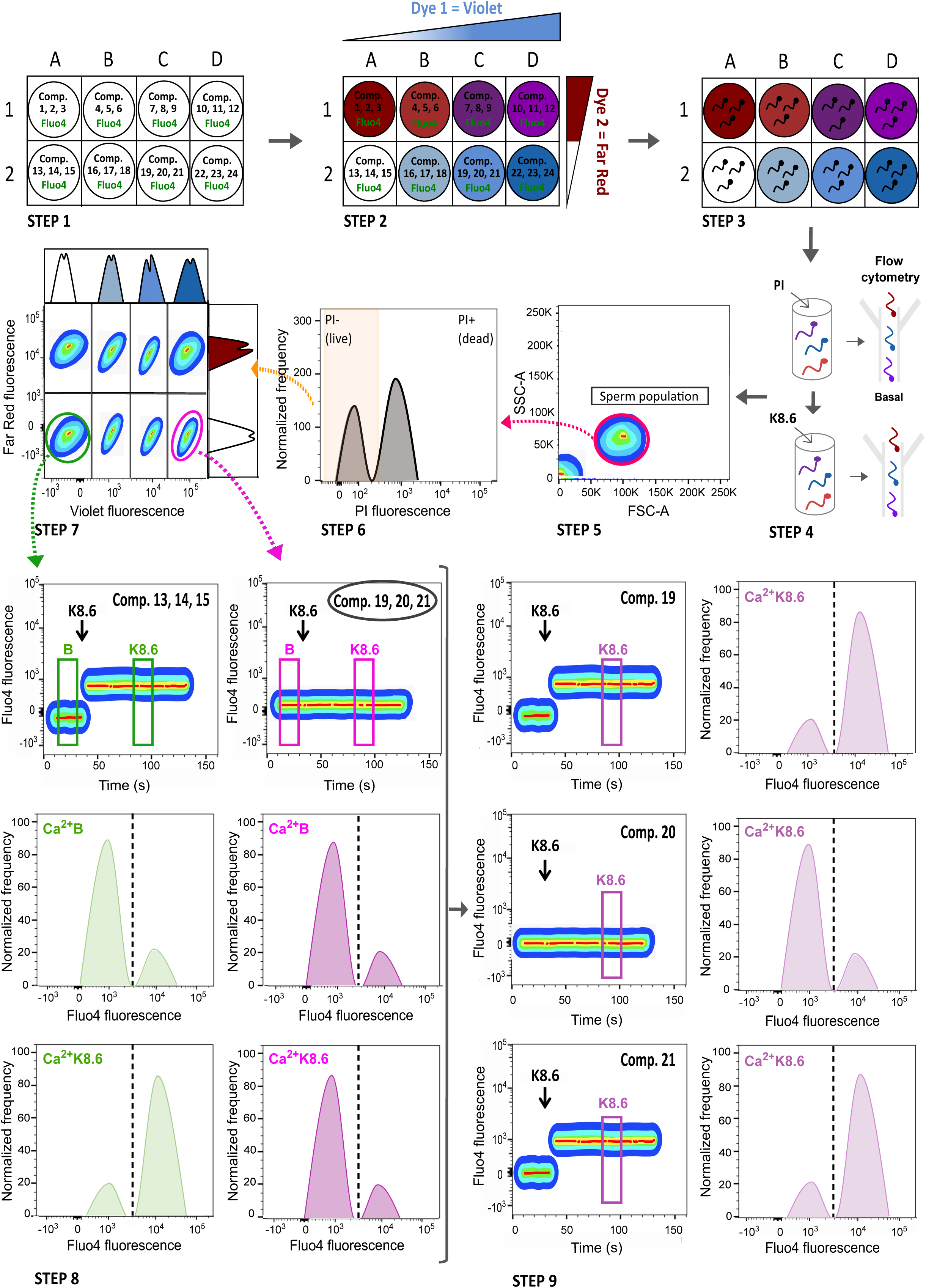
FCB–K8.6 screening workflow for CatSper activity. (1) A 96-well plate is prepared with three test compounds per well and the Ca^2+^ indicator Fluo-4. (2) Each well receives a unique CellTrace™ Violet/Far Red dye combination for barcoding. (3) Sperm are added to each well, and incubated 20 min with compounds and dyes. (4) Samples are pooled, stained with PI, and analyzed by flow cytometry. Basal [Ca^2+^]ᵢ is recorded for 30 s, then the NC medium is replaced with K8.6 medium to induce CatSper activation, followed by a 90 s recording. (5–7) Live (PI^-^) sperm are gated and decoded by CellTrace™ Violet vs. Far Red signatures. (8) Fluo-4 fluorescence before (B) and after K8.6 identifies compound mixtures that inhibit CatSper (no [Ca^2+^]ᵢ rise). (9) Compounds from inhibitory wells are then tested individually.

As shown in Figure 3**, step 8**, control pools (green histograms) displayed the expected increase in Fluo-4 signal upon K8.6 stimulation, reflecting the rise in [Ca^2+^]ᵢ, whereas pools containing active CatSper inhibitors showed a blunted response (pink histograms). Pools showing inhibition were subsequently analyzed individually to identify the active compound responsible (Figure 3**, step 9**). In this schematic example, compound 20 shows inhibitor effect on the CatSper-mediated increase in Ca^2+^ upon K8.6 stimulation.

Figure 4A shows a representative 4×2 matrix in which different sperm populations can be resolved according to the concentration of barcoding dyes. The position of each treatment within the matrix is shown schematically as a double-entry table, defined by its CellTrace™ Violet vs. CellTrace™ Far Red fluorescence signature. For instance, in position V4–Fr1, a known CatSper inhibitor (20 μM HC-056456) was pooled with two TRPV1 antagonists (5 μM CZ and 250 nM BCTC), which are not expected to affect CatSper activity. As a negative control, DMSO (vehicle) was included at position V1–Fr1.

**Figure 4.**
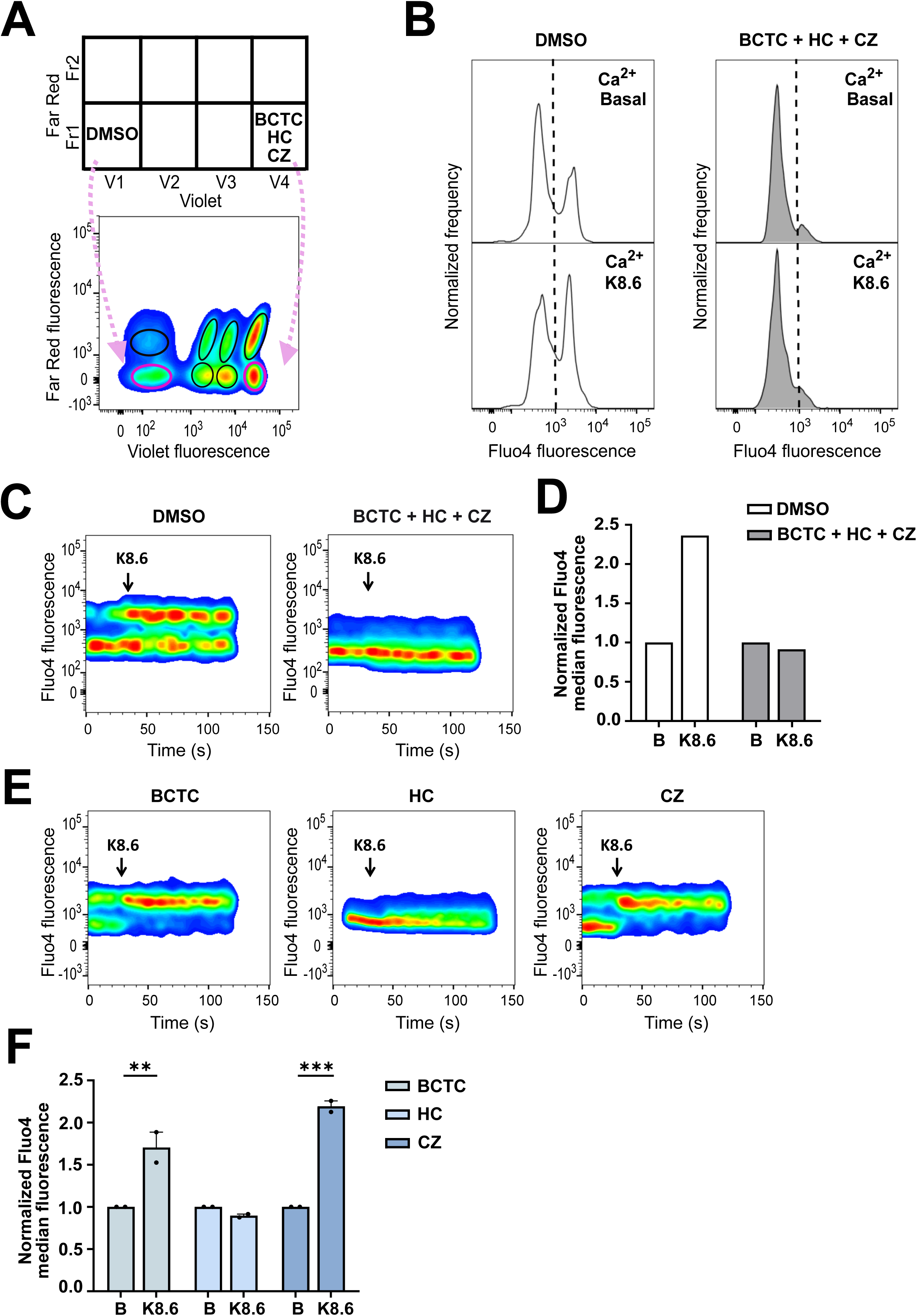
Application of the FCB–K8.6 method to identify CatSper inhibitors. **(A)** Schematic of the 4×2 matrix containing one well with a negative control (DMSO) and one well with a positive control: compounds combinations (BCTC, HC, CZ), with HC a well-known CatSper inhibitor. Barcoding dye concentrations (Violet: V1 = 0.025 μM; V2 = 1.5 μM; V3 = 5 μM; V4 = 20 μM, and Far Red: Fr1 = 0.001 μM; Fr2 = 0.4 μM) are shown in a double-entry table. A representative CellTrace™ Violet vs. Far Red dot plot shows the eight sperm subpopulations; pink arrows indicate two marked populations. **(B)** Representative histograms of Fluo-4 fluorescence before (basal [Ca^2+^]i) and after K8.6 stimulation for negative control and CatSper inhibitor-treated sperm. **(C)** Time-course plots of Fluo-4 fluorescence showing that K8.6 induced a [Ca^2+^]ᵢ rise only in vehicle-treated cells. **(D)** Median Fluo-4 fluorescence before (B) and after K8.6 stimulation normalized to the basal condition. **(E–F)** Analysis of individual compounds shows that only HC blocked the K8.6-induced [Ca^2+^]ᵢ response (n=2). ** p < 0.01; *** p < 0.001 indicates statistical difference.

After selecting each barcoded population, Fluo-4 fluorescence was monitored over time (Figure 4B**–C**), and median fluorescence values before and after K8.6 stimulation were quantified (Figure 4D). As expected, the presence of HC-056456 prevented the K8.6-induced rise in [Ca^2+^]ᵢ (Figure 4B**–D****, pool**), whereas DMSO-treated sperm displayed the characteristic increase in fluorescence (Figure 4B**–D****, vehicle**). Subsequent analysis of each compound in the inhibitory pool (Figure 4E**–F**) confirmed that the effect was specifically due to HC-056456, as neither CZ nor BCTC altered the Ca^2+^ response.

Overall, this workflow establishes a robust and scalable method for the functional evaluation of CatSper activity and the possible identification of novel inhibitors, integrating single-cell Ca^2+^ measurement, FCB, and compound pooling into a single high-throughput assay.

## Discussion

CatSper has long been considered a highly promising target for the development of a non-hormonal male contraceptive as is exclusively expressed in sperm and is indispensable for male fertility [30,33,42]. However, progress toward identifying CatSper-targeting compounds has been limited, largely due to the lack of scalable, robust, and physiologically relevant assays to evaluate channel function and screen for potential inhibitors.

Fluorescence-based assays that infer CatSper activity by monitoring changes in Em have proven to be highly informative and, particularly when adapted to flow cytometry, enable single-cell resolution [21,22,30]. However, because Em can also be shaped by the activity of other ion channels or electrogenic transporters, Em-based readouts may not exclusively reflect CatSper function. For this reason, complementary strategies that directly monitor Ca^2+^ entry are essential. Here, we implemented a Ca^2+^-based flow cytometry assay that provides a direct, sensitive, and scalable measurement of CatSper-mediated Ca^2+^ influx. Importantly, this assay was validated using CatSper1 KO sperm, which failed to exhibit the K8.6-induced rise in [Ca^2+^]ᵢ. While we cannot completely exclude minor contributions from other Ca^2+^-conducting pathways, the markedly reduced signal in CatSper1 KO sperm indicates that such contributions are not sufficient to impact the interpretation of the results obtained with this methodology.

Stimulation of CatSper by high-K^+^/high-pH medium (K8.6), which simultaneously depolarizes the membrane and increases intracellular pH, effectively promotes channel opening because alkalinization shifts the voltage dependence of CatSper toward more negative potentials, enabling Ca^2+^ influx [9,19]. This pH-dependent gating is mainly regulated by the CATSPER ζ –EFCAB9 complex, where EFCAB9 acts as a Ca^2+^-sensitive EF-hand protein that adjusts channel responsiveness to intracellular pH. In Efcab9 KO sperm, that simultaneously lack CatSper ζ, CatSper channels are present but display markedly diminished activation upon alkalinization, indicating that this complex functions as a dual pH/ Ca^2+^ sensor that ensures appropriate channel opening [14]. Notably, intracellular alkalinization facilitates CatSper activation in both mice and human sperm [19,20], unlike other modulatory mechanisms (e.g., progesterone or prostaglandins) which are human-specific [43,44]. This conservation of pH-dependent gating underscores the relevance of alkalinization-based assays, such as the one implemented here, and supports their adaptability across species.

The compatibility of FCB with Fluo-4 labeling and viability discrimination demonstrates that this strategy can be applied to live sperm without altering Ca^2+^ measurements. Moreover, because flow cytometry allows the simultaneous detection of several fluorescent probes, as long as their emission spectra are appropriately separated, including through multispectral flow cytometry, this platform can be readily expanded to assess multiple capacitation-associated parameters in parallel. In this way, intracellular pH, Na^+^ dynamics, reactive oxygen species, mitochondrial activity, or acrosomal status can be measured alongside CatSper-dependent Ca^2+^ influx, substantially broadening the analytical power and applicability of this approach.

In addition, we introduced a compound pooling strategy to increase screening efficiency. Pooling significantly decreases the number of assays required to survey large compound libraries, an advantage that is particularly relevant during early hit-finding phases. However, pooling designs must consider potential compound–compound interactions, including additive, synergistic, or antagonistic effects. Synergy may generate false positives, whereas antagonism may mask inhibitory effects and result in false negatives. Moreover, chemical incompatibility within pools may lead to aggregation or degradation [41]. For these reasons, pools showing inhibitory activity must be deconvoluted through secondary testing of individual compounds. Despite these considerations, pooling remains a highly effective strategy for high-throughput phenotypic screening when used in combination with appropriate validation steps. In our system, the initial pooling approach enabled the rapid identification of inhibitory mixtures, followed by straightforward deconvolution to pinpoint the active molecule.

In conclusion, we establish a robust, scalable, and high-throughput assay for evaluating CatSper activity in live sperm. By combining K8.6-induced Ca^2+^ entry, flow cytometry, FCB, and compound pooling, this platform provides a powerful tool for discovering novel CatSper inhibitors and advancing the development of non-hormonal male contraceptive candidates. Beyond its application for drug discovery, this method offers a versatile framework to explore fundamental aspects of sperm physiology with single-cell resolution.

---

Supp. Table 1: Key drugs table.

## Acknowledgments

We would like to thank all the members of our laboratory for their insightful comments. We gratefully acknowledge the financial support provided by the Williams, Bigand and Rene Baron Foundations, as well as the *Sociedad Argentina de Biología* (Eduardo Charreau award to GML).

## Author contribution

LRGO, LJSE and MJ performed experiments; LRGO analyzed the data; LRGO and GML prepared the manuscript with contributions of all other authors; GML and MGB designed the study. GML and MGB acquired the funding.

