## Supplementary material for "A high-throughput flow cytometry assay to evaluate CatSper-mediated Ca^2+^ influx triggered by high-K^+^/high-pH stimulation in mouse sperm": Suppl Table 1

| Reagent | Company | Catalog # | Concentration used | Vehicle |
|---|---|---|---|---|
| Fluo-4 AM | Invitrogen, Thermo Fisher Scientific (USA) | F14201 | 1 μM | DMSO |
| Pluronic acid | Invitrogen, Thermo Fisher Scientific (USA) | F14201 | 0.02% | DMSO |
| CellTrace™ Violet | Invitrogen, Thermo Fisher Scientific (USA) | C34557 | 0.025 μM, 1.5 μM, 5 μM, 20 μM | DMSO |
| CellTrace™ FarRed | Life Technologies, Thermo Fisher Scientific (USA) | C34564 | 0.001 μM, 0.4 μM | DMSO |
| HC-056456 | MedKoo Biosciences (USA) | 531948 | 20 μM | DMSO |
| RU1968 | Kindly provided by Dr. Timo Strünker | - | 10 μM | DMSO |
| [N-(4-tert-butylphenyl)-4-(3-chloropyridin-2-yl)piperazine-1-carboxamide] (BCTC) | Tocris (United Kingdom) | 3875 | 250 nM | DMSO |
| 5-(N,N-hexamethylene)-amiloride (HMA) | Cayman Chemical (USA) | 29788 | 10 μM | DMSO |
| Capsazepine (CZ) | Cayman Chemical (USA) | C191 | 5 μM | DMSO |
| Propidium iodide (PI) | Santa Cruz Biotechnology (USA) | SC3541 | 2 ng/μL | $H_2O$ |
